# SARS-CoV-2 mutations and where to find them: An *in silico* perspective of structural changes and antigenicity of the Spike protein

**DOI:** 10.1101/2020.05.21.108563

**Authors:** Ricardo Lemes Gonçalves, Túlio César Rodrigues Leite, Bruna de Paula Dias, Camila Carla da Silva Caetano, Ana Clara Gomes de Souza, Ubiratan da Silva Batista, Camila Cavadas Barbosa, Arturo Reyes-Sandoval, Luiz Felipe Leomil Coelho, Breno de Mello Silva

## Abstract

The recent emergence of a novel coronavirus (SARS-CoV-2) is causing a severe global health threat characterized by severe acute respiratory syndrome (Covid-19). At the moment, there is no specific treatment for this disease, and vaccines are still under development. The structural protein Spike is essential for virus infection and has been used as the main target for vaccine and serological diagnosis test development. We analysed 2363 sequences of the Spike protein from SARS-CoV-2 isolates and identified variability in 44 amino acid residues and their worldwide distribution in all continents. We used the three-dimensional structure of the homo-trimer model to predict conformational epitopes of B-cell, and sequence of Spike protein Wuhan-Hu-1 to predict linear epitopes of T-Cytotoxic and T-Helper cells. We identified 45 epitopes with amino acid variations. Finally, we showed the distribution of mutations within the epitopes. Our findings can help researches to identify more efficient strategies for the development of vaccines, therapies, and serological diagnostic tests based on the Spike protein of Sars-Cov-2.

## Introduction

The SARS-CoV-2, a novel coronavirus, is highly transmissible, leading to high infection rates and human mortality around the world, turning the disease caused by this virus (COVID-2019) a huge public health concern (Li et al., 2020). Up to now, SARS-CoV-2 has been spreading in several continents and causing more than 4,629,000 confirmed cases, with mortality rates of 6,74% (World Health Organization, 2020 – access at 18/05/2020).

Coronaviruses have the largest genome among RNA viruses (26 to 32 kilobases in length) with 14 ORFs encoding 27 proteins. At 5’ end of the genome, there are the 1ab and 1a ORFs, which encode 16 mature non-structural proteins (nsp1 to nsp16). These proteins play crucial functions during viral RNA replication and transcription^1^. At 3’ end of the genome, there are genes encoding four structural proteins: Spike (S), Envelope (E), Membrane (M), and Nucleoprotein (N) as well as genes for 8 accessory proteins named 3a, 3b, p6, 7a, 8b, 9b, and orf14^2^.

Although recent findings suggest that SARS-CoV-2 has stronger transmissibility when compared to SARS-CoV, the molecular mechanisms responsible for this difference remain unclear^3^. However, among the structural proteins, the Spike glycoprotein that is present as a homo-trimer on the coronaviruses surface, has been pointed as the most important factor responsible for this stronger transmissibility since this protein is able to bind cell receptors. The S protein has two subunits named S1 and S2. The S1 subunit is responsible for binding to the host cell receptor. It has a signal peptide, an N-terminal domain (NTD) and a receptor-binding domain (RBD). The S2 subunit is responsible for fusion of the viral and cellular membranes and consists of a conserved fusion peptide (FP), two hepta-repeats (HR1 and HR2), a transmembrane (TM) and a cytoplasmic (CP) domains^4^. Structural analysis of the receptor-binding domain (RBD) of SARS-CoV-2 Spike protein and the human receptor angiotensin-converting enzyme 2 (ACE2) revealed that the RBD can induce the spillover to other animals as well as human-to-human transmissions^5,6^. This protein has been shown as a key factor for coronaviruses entry into cells, as well as a target for neutralizing antibodies, due to its role in binding to cellular receptors and fusion of viral and cellular membranes ^4,7,8^.

The RBD of SARS-CoV S1 domain undergoes conformational changes that hide or expose the determinants of receptor binding^6,9,10^. Then, upon RBD binding to cellular receptors, the Spike protein is cleaved by proteases and the signal peptide is released. This cleavage triggers conformational changes in the S1 and S2 subunits, leading to the exposure of the fusion loop and its interaction into target cell membrane. This fact turns this domain a target to virus neutralization by monoclonal antibodies. Thus, conformational changes of the Spike protein are a necessary step to viral membrane fusion, and it allows the entry of viral nucleocapsids into the host cell to initiate replication^7,9,11,12^. Additionally, the S glycoprotein of SARS-CoV-2 has a furin-like cleavage site at the S1/S2subunits, which highlights the essential differences in Spike protein between SARS-CoV and SARS-CoV-2. Thus, this mechanism is proving to be a potential target for vaccines, therapeutic approaches, and diagnosis for coronaviruses^13,14^.

Since the Spike protein has a crucial role in the initial steps of SARS-CoV-2 replication, research studies have been focusing on its structure, function, and antigenicity to gain a better understanding ofthe Spike protein^14,15^. A bioinformatics analysis has shown that the S2 subunit of SARS-CoV-2 is highly conserved and shares 99% identity with those of the two bat SARS-like CoVs (SL-CoV ZXC21 and ZC45) and human SARS-CoV^8^. In addition, the tri-dimensional model of Spike protein structure has recently been published, showing that the SARS-CoV-2 Spike protein has more affinity for binding ACE2 than the SARS-CoV Spike protein^16^.

However, some studies have shown that cellular receptors used to viral attachment and entry can vary among host species of different coronaviruses ^9,14,17,18^. A recent study reported that SARS-CoV-2 is most closely related to the bat SARS-CoV RaTG13 that forms a distinct lineage of SARS-CoVs, and their Spike glycoproteins share 98% amino acid sequence identity^19^. Despite the amino acid level appears to be similar, there are important differences in Spike protein between these viruses that might explain some viral differences regarding the pathogenicity ^14^. Therefore, more studies are needed to understand how those differences can change SARS-CoV-2 functionality.

Data about the genomic variability of the SARS-CoV-2, especially on the region encoding the Spike protein, could provide important support for the accuracy of structural predictions. In this present study, we analysed 2363 sequences of the Spike protein in order to study the frequency of mutations on the protein domains and motifs and to identify potential epitopes on this protein. Additionally, we also determined the epitope variability of the Spike protein circulating in all continents. This work can be used to support long-term studies to identify Spike protein mutations emergence and understand how it can affect vaccine trials and serological diagnosis.

## Materials and methods

### Sequence retrieval and Structural analysis

Only complete genomes of SARS-CoV-2 were collected in the GISAID (https://www.gisaid.org/). Complete sequences of Spike protein of SARS-CoV-2 were obtained from NCBI (https://www.ncbi.nlm.nih.gov/) and ViPR (https://www.viprbrc.org). All sequences are aligned using MEGA-X^20^ software with MUSCLE algorithm^21^.

The homo-trimer model was obtained through the Robetta server^22^ (http://robetta.bakerlab.org/) referenced by the partial crystal of the “closed-state” of SARS-CoV-2 Spike protein6VXX^23^, representing about 77% of the actual structure (https://www.rcsb.org/structure/6VXX). Subsequently, the structure of model was subjected to geometry optimization steps by the Discovery Studios software^24^, where was considered eleven outlier residues from the favorable/acceptable regions in the Ramachandran plot: 59PHE^A,B,C^, 62VAL^A,B,C^, 365TYR^A,B,C^, 544ASN^B^ and 744GLY^B^. So that all residues with a radius distance of 4A from each of the outliers was also considered in the optimization steps.

### T and B cell epitope prediction

All obtained sequences were used to predict T and B cell epitopes. The Wuhan-Hu-1 Spike protein of reference sequence NC_045512.2^25^ (https://www.ncbi.nlm.nih.gov/protein/1796318598) was used as reference for the epitope prediction. The NetCTL1.2^26^ (http://www.cbs.dtu.dk/services/NetCTL/) was used to predict MHC-I binding epitopes. The MHC class I was considered for the prediction of epitopes for cytotoxic T cells through artificial neural networks, using the standard set of Weight on C terminal cleavage score (0.15), Weight on TAP (Transporter Associated with Antigen Processing) efficiency matrix (0.05) and Threshold for epitope identification (0.75). NetMHCpan 4.0^27^ (http://www.cbs.dtu.dk/services/NetMHCpan/),was also used to predict MHC class I epitopes for cytotoxic T cells. Peptides with mers of 8-11 was pointed through artificial neural networks, with the threshold of <2% better in the binding score rank. The NetMHCII 2.3^27^ (http://www.cbs.dtu.dk/services/NetMHCII/), was used to prediction epitopes for T Helper cells with 15 mers. The *loci*HLA-DR was used, with standard threshold (<2% better in the binding rank affinity) to identify the peptides that best bound to MHCII.

Linear epitopes for B cells of different sizes were predicted using BepiPred-2.0 (http://www.cbs.dtu.dk/services/BepiPred/)^28^. The standard threshold of 0.5 was used to ensure the better Specificity/Sensitivity ratio of the epitope. Finally, the conformational epitopes for B cells were predicted using the validated model of the three-dimensional structure of the spike protein homo-trimer through two web servers (DiscoTope and hroughElliPro). DiscoTope (http://tools.iedb.org/discotope/)^29^ was used with the threshold for specify epitope identification (−3.7). The prediction of conformational epitopes using ThroughElliPro^30^(http://tools.iedb.org/ellipro/) was made with the maximum score threshold of 0.5 and the maximum distance for ligation of 6 angstroms. Only those epitopes located in the “extracellular” region of the protein (13-1213) were considered. The identified epitopes were visualized on Spike protein tri-dimensional model using the software Visual Molecular Dynamics^31^ (figures 1 to 3) or Discovery Studios (supplementary figure 2) ^32^. The image processing for figures 1 and 3 was done using the software Visual Molecular Dynamics^31^. The flowchart of the three main stages of the study is on figure 1.

**Figure 1.**
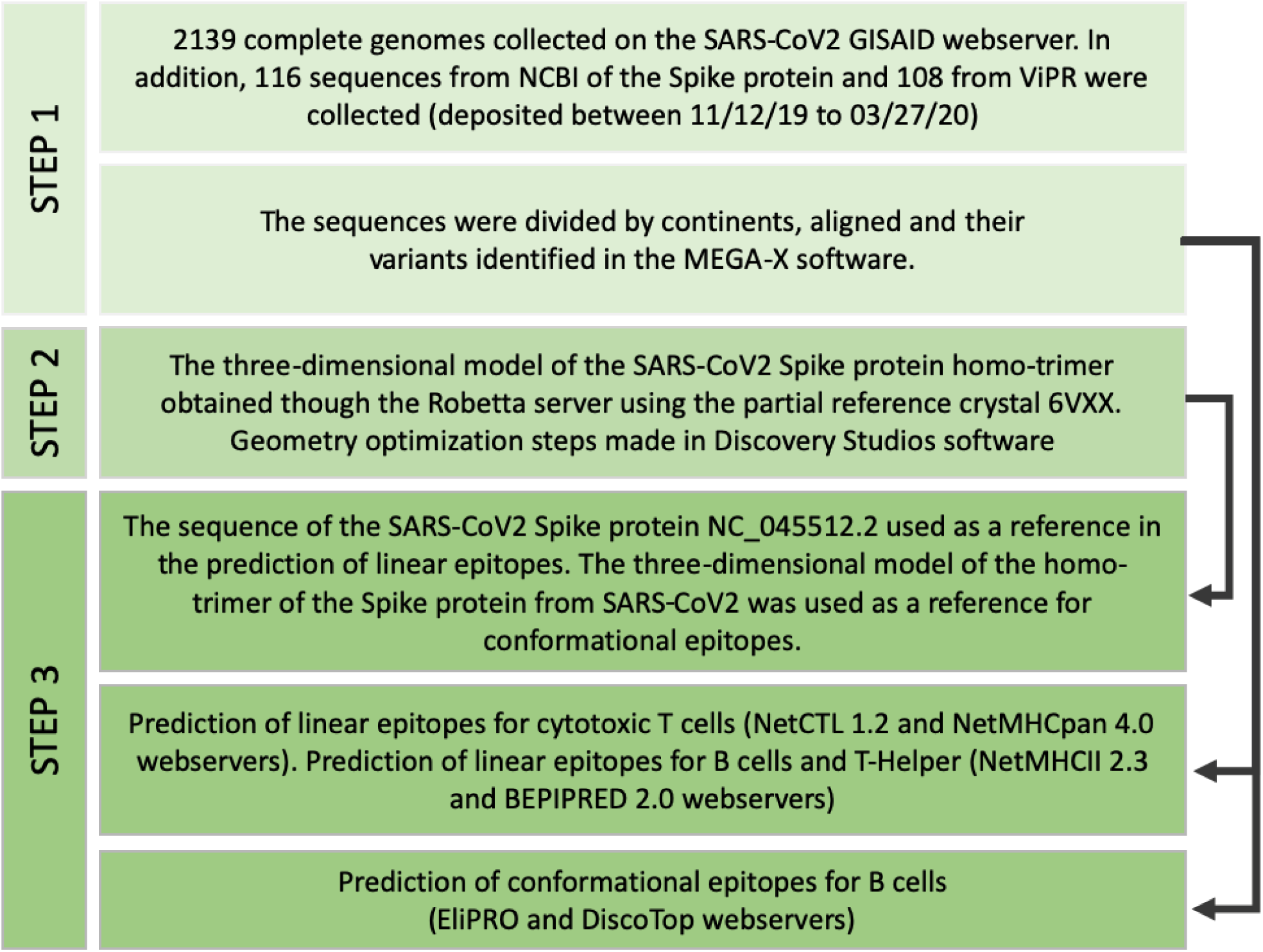
The general outline of the methodology. Flowchart of the three main stages of the study.

**Figure 2.**
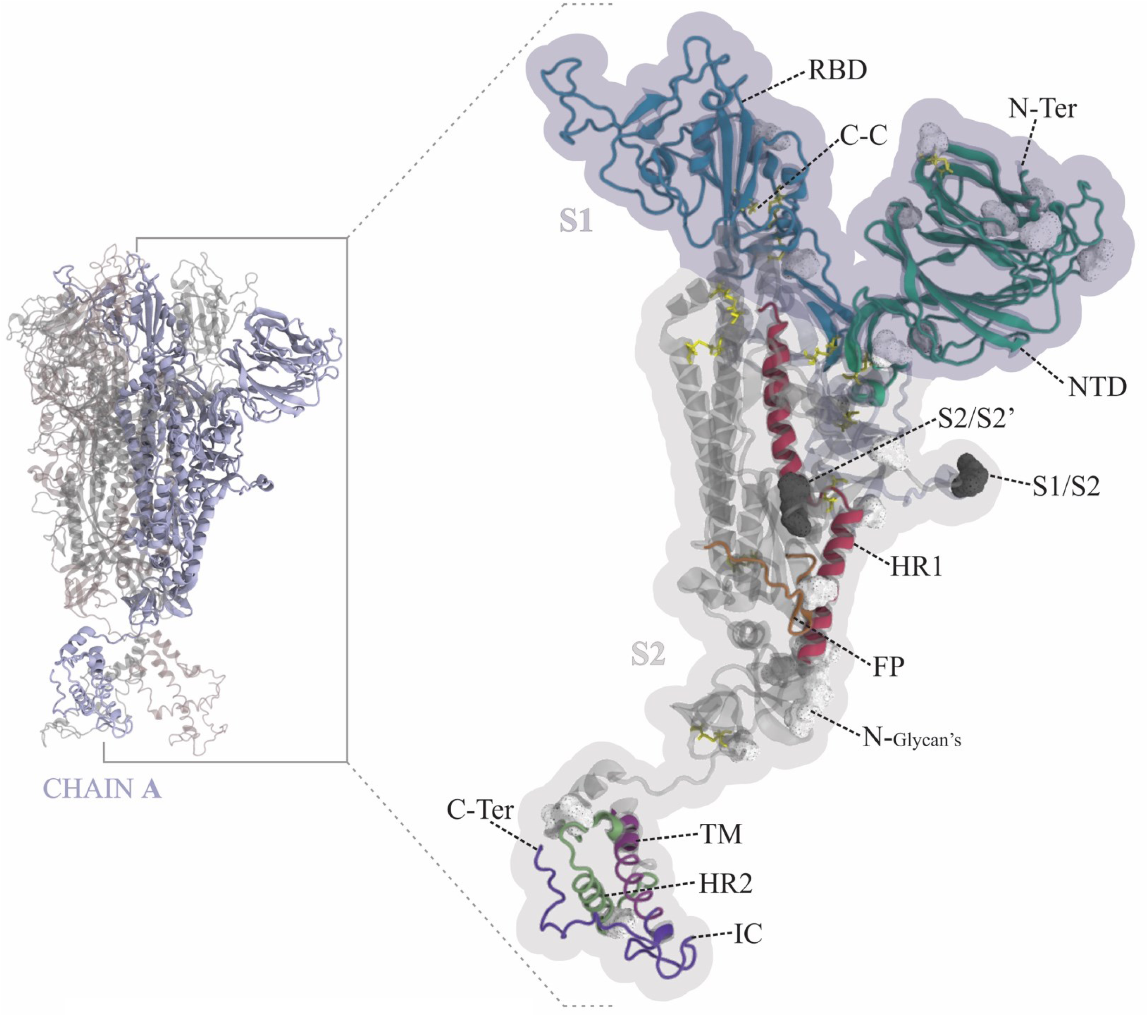
Structural mapping of SARS-CoV-2 Spike protein: On the left side, the homo-trimer is showed as a newcartoon. The B and C chains are represented in gray and the A chain in blue. On the right side, the monomer is also showed as a newcartoon with the representation of domains, subdomains, motifs, N-Glycan’s, and disulfide bonds. The blue and gray shadows above structure represent the portions of S1 and S2 domains, respectively. (See video 1 in supplemental material)

**Figure 3.**
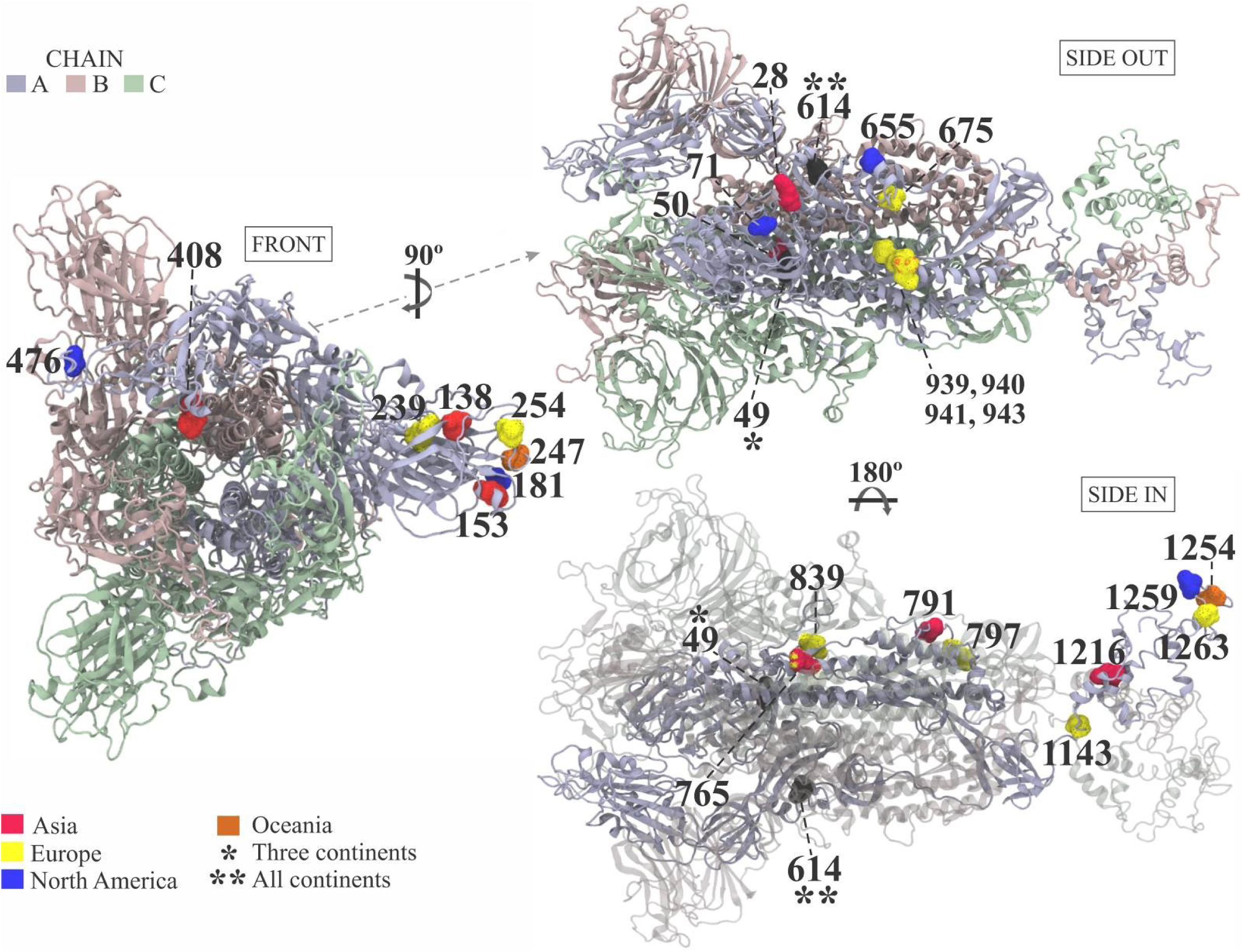
SARS-CoV-2 Spike protein variant amino acid residues: The homo-trimer structure is represented in newcartoon. The positions with the mutations among residues of non-homologous physico-chemical characteristics are represented on the surface in the A chain (light blue) only. In the “SIDE IN” perspective, the B and C chains are represented in transparent newcartoon. The colors of each residue are represented by the continent where the variations were identified.

## Results

### Structural modeling

The monomeric Spike protein model was represented according to their respective domains, disulfide bridges, and glycosylation points indicated by Expasy’s P0DTC2 annotations reports (https://viralzone.expasy.org/resources/Coronav/P0DTC2.txt) (figure 2 and video 1 in supplemental material). The FP region, reported from Expasy’s, has a little difference than that is reported for Sars-Cov. The three-dimensional Spike homo-trimer model showed a clash score of 1.13 and a molprobity (http://molprobity.biochem.duke.edu/) score of 1.05 corresponding to the 99th and 100th percentiles, respectively, when compared to high-resolution 3D structures. The model revealed 99.9% of its residues in acceptable/favorable regions on the Ramachandram Plot (http://molprobity.biochem.duke.edu/), where only 59PHE residues from chains A, B and C continue as an outlier (supplementary figure 1). After ensuring the model quality, the homo-trimer 3D structure was used to the structural/antigenic SARS-CoV-2 Spike protein variability mapping.

### Variations of protein Spike

After Spike protein sequences retrieval and comparison, 856 amino acid variations were identified in 32.8% of sequences. Europe and South America showed the highest variation rates among worldwide sequences, showing variation on 47.4% and 44.1% of sequences, respectively. (data not showed). Sequence comparison analysis identified 42 amino acid residues with variation that occurred at least twice on SARS-Cov-2 sequences. Twenty three variations was mapped in the S1 and 19 in S2 domain (table 1). Some of these variations (28) are represented by residues with non-homologous physical-chemical characteristics in their respective continents of occurrence (figure 3).

**Table 1.**
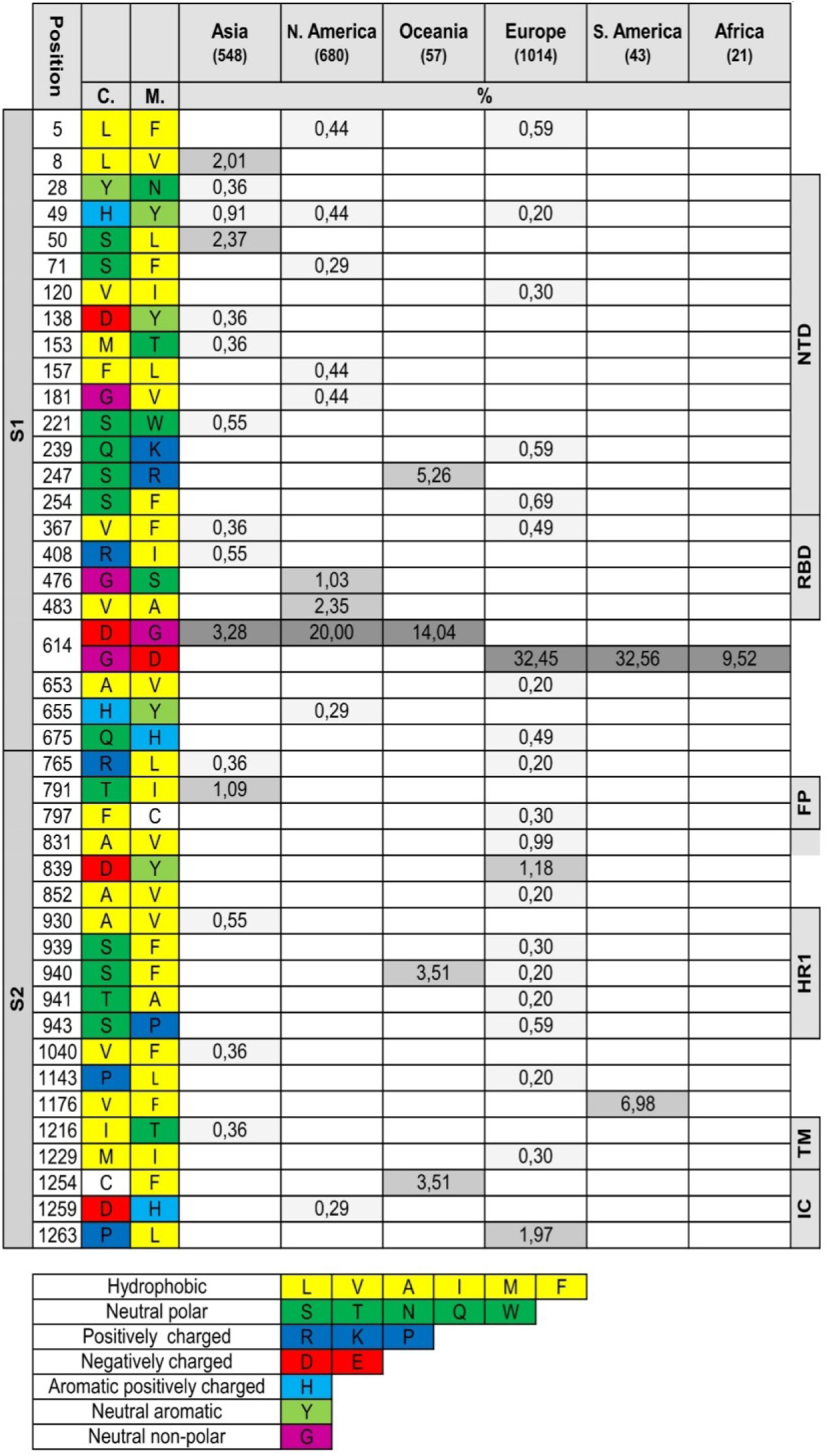
SARS-CoV-2 Spike protein amino acid variations: All variations are identified by their position in the polypeptide chain, as well as the physical-chemical characteristic of the consensus (C.) / Variant (M.) Residue. The residues were pooled on continents from which the samples were isolated. Protein domains are indicated on the left and the subdomains and motifs on the right. Mutations that occur at frequencies< 1% are indicated by light gray, > 1% mutations by gray and 614 variations by dark gray.

### Epitopes prediction of protein Spike

In the present analysis, it was predicted 282 epitopes: 95 for T-cytotoxic cells, 135 for T-Helper cells and 52 for B cells (30 linear and 22 conformational epitopes). All epitopes are shown in supplementary table 1. Only those variations whose showed amino acid changes to residues with non-homologous physical-chemical characteristics was considered and represented in table 2. As a result, there are 11 predicted epitopes for T-Cytotoxic cells, 16 for T-Helper cells, 18 for B cells (10 linear and 8 structural epitopes). Forty-five epitopes with mapped variations were represented in the 3D model of the SARS-CoV-2 protein Spike structure (figure 4). The S1 domain gathers the majority counts (33) while the NTD region showed 20 epitopes, of which 7 were predicted for B cells (table 2).

**Table 2.**
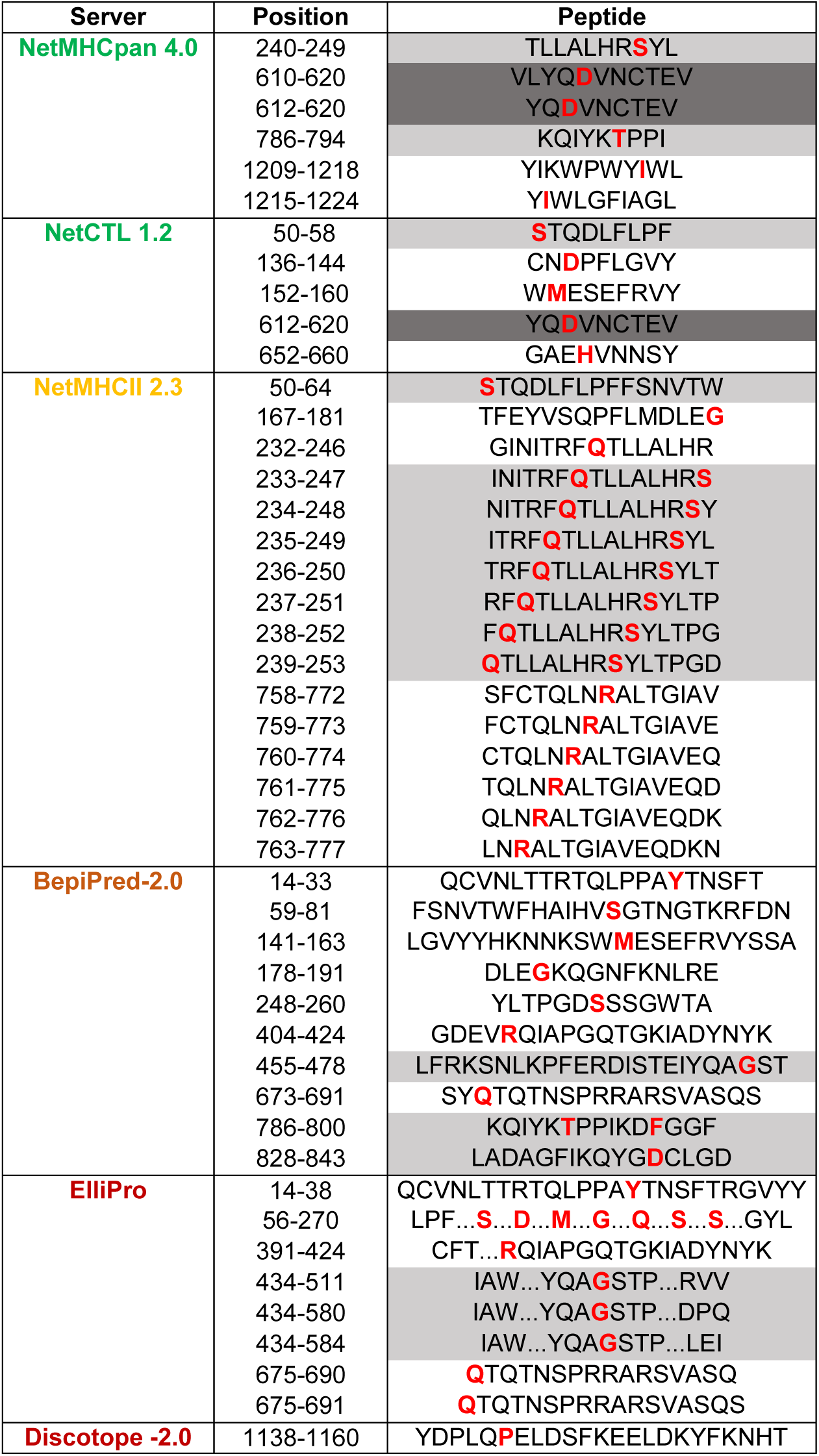
SARS-CoV-2 protein Spike identified epitopes with amino acid variants: The servers in green represent the T-cytotoxic cells and yellow for T-Helper cells predictions. Antigenic prediction servers of B cells are shown in red and orange colors, for conformational and linear epitopes, respectively. All epitopes are identified by their positions in the polypeptide chain, as well as by their amino acid sequences. A highlighted letter in red represents the variation present in each epitope. The highlight in light gray represents the presence of mutations with frequencies on the coding region > 1% and the epitopes with the mutations in residue 614 are represented by dark gray.

**Figure 4.**
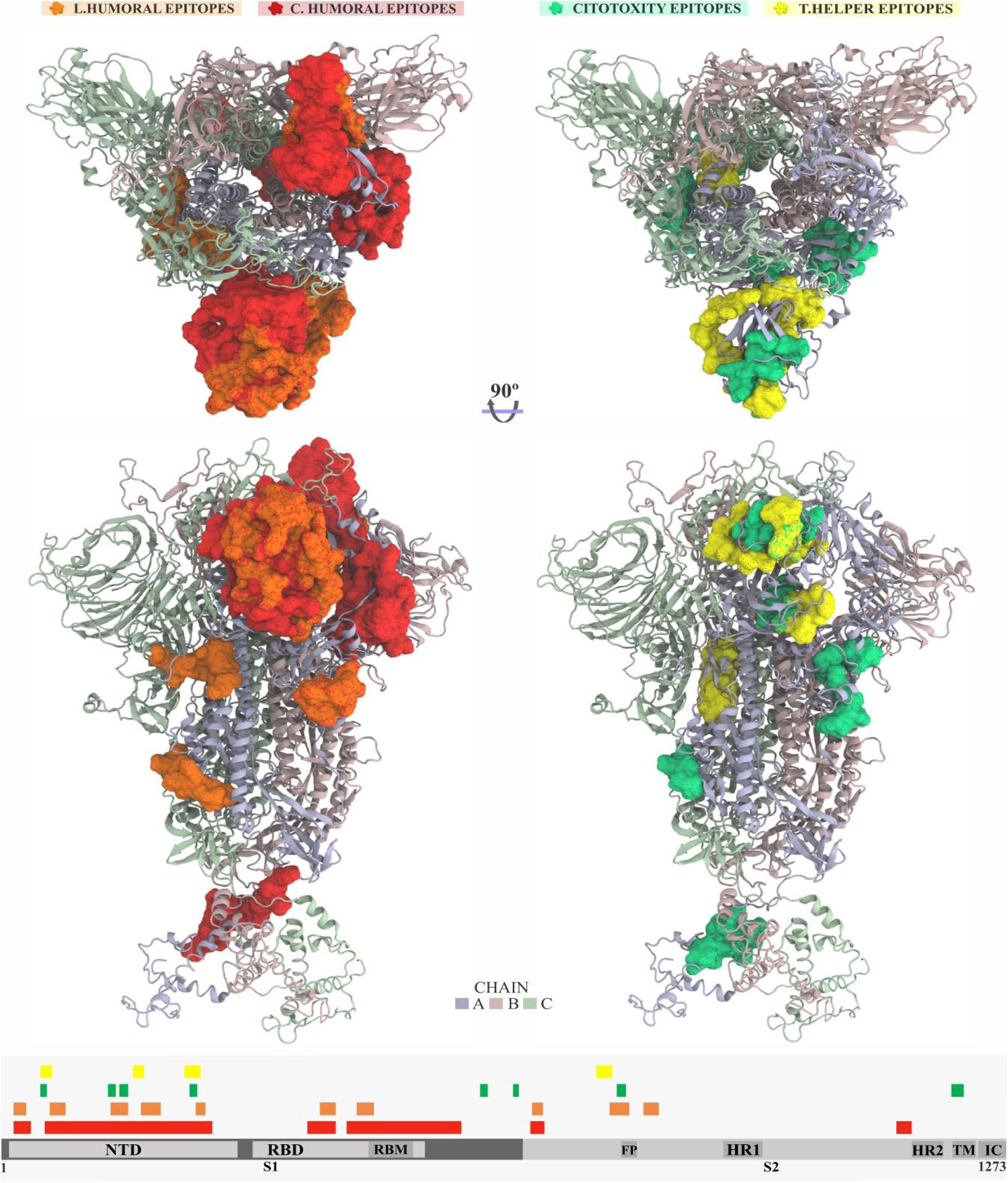
Mapping of epitopes with variants on SARS-CoV-2 Spike protein: The homo-trimer structure was represented in newcartoon. Epitopes are represented by their surfaces in the A chain, in colors that represent the type of prediction as highlighted at the top of the figure. The histogram below the structures represents the Spike protein polypeptide chain with its domains, subdomains, motifs and epitopes proportionally represented in relation to the primary sequence of the proteins.

## Discussion

### Amino acids variations

Our findings suggest that the majority of residue variations arose punctually on their respective continents, with the exception of position 614, which occurs in all continents. The 614-residue variation has the most expressive frequency and has been reported by recent published/preprinted studies ^33–35^. The H49Y variation has been identified in Asia, Europe, and North America. In contrast, L5F, V367F, R765L, and S940F variations, with the exception the last, were observed in at least two continents with <1% frequency. (table 1).

Overall, the largest number of variants was found in S1 protein subunit, with 10 variant positions found in Asia, 9 in Europe, 9 in North America, and 2 in Oceania (table 1). South America and Africa did not show any other variation in the S1 domain besides that identified in residue 614, probably due to the few numbers of analyzed genomes from these regions. The sequences from Europe showed a higher number of variant residues in the S2 domain (12) in comparison to the average from the other continents (1.8). However, the S1 domain is the region more propense to appear new variations due its most prominent variability between coronaviruses species^36^.

Due to the short period since its dissemination, it is difficult to find a specific variation pattern in Spike protein of SARS-Cov-2 on the continents. But even so, the variant position D1259H in S2 domain was seen in North America sequences but not in Asia and Europe (table 1). Future works with a higher number of high-quality sequences are needed to better understand the possible heterogeneous distribution of South/Central Americas, Africa and Oceania amino acid variations on these regions.

### Mapping of variations

#### S1 DOMAIN

Our analysis showed that the NTD domain has many variations (figure 3), where changes on teen residues were identified. However, only two residues variations showed frequency above 1% (S50L and S247R). The S50L residue variation was found in 2.37% of sequences from Asia and is internally located in the homo-trimer structure. A recent study demonstrated that the L50 residue found in SARS-Cov / SARS-like CoV’s was replaced by S50 in SARS-CoV-2 ^2^. However, some SARS-Cov-2 sequences remain with the L50 residue. Another particular residue variation (S247) showed 5.26% frequency on Oceania. This residue is located at the peripheral end of the NTD portion and was reported in a recent preprint study that five variations of Spike protein may be related to variation of replication of SARS-CoV-2 in Vero-E6 cells^37^.

In the RBD region, four residues variations were identified (table 1). A recent in silico preprint study shows the variation V367F as a probable factor to the high increase of binding affinity with the ACE2 receptor^40^. The V483A variation, found only in North American sequences, are co-located in the binding region of the main residues involved in the interaction of the SARS-Cov Spike protein with the ACE2 receptor ^38,39^. The R408I variation was identified only in the Asian continent and it was reported in a recent in silico preprint study as a probable factor of decreased binding affinity with the ACE2 receptor^40^ (figure 3-front & table 1). Additionally, the arginine at position 408 in RBD core region has already been cited as an important point of interaction with N-glycan, for both SARS and SARS-CoV-2^41,42^. Thus, future investigations should be done in order to evaluate if these changes on Spike protein residues could be correlated to functional changes.

A second G476S variation observed in the North American genomic sequences was found in peripheral and accessible region of the structure (figure1-front) and is also co-located with the region of the main interaction residues between the SARS-CoV-2 protein Spike and the ACE2 receptor ^43^. This was also observed inSARS-CoV^39^. As mentioned before, the highest frequency mutation rate was identified at position 614 in almost all continents (except Central America). This variation could be used to group the continents into either D or G predominantly variations. The group D includes Asia, North America, and Oceania while group G includes Europe, South America, and Africa (table 1). However, although residue D614 in SARS and SARS-CoV-2 reference sequences indicates a positive selection of G614 mutation in those continents, there is no evidence of how this mutation frequency could impact virus functionality.

Coronavirus SARS-CoV-2 sequencing analysis from diverse continents demonstrates that the genes encoding the Spike proteins undergo diverse and frequent mutations^44^. First, it was identified a mutation (241C>T), which was developed gradually, and reported with three other co-mutations (3037C>T, 14408C>T, and 23403A>G)^33^. These mutations culminated on amino acid variations in Spike protein, nsp3, and RNA primase, which are responsible for RNA replication^33^. Moreover, all associated-mutations observed were prevalent in Europe isolates, giving insights for SARS-CoV-2 severity in this region. Therefore, the SNP mutation (23403A>G/D614G) in Spike protein D614G, was pointed out by Yin, C.^33^. This residue is near to this furin recognition site (S1/S2 region) of SARS and could affect enzyme activity. However, the distance between alpha carbons of residues 614 and 685 turns the cleavage site for SARS-CoV-2 (S1/S2) greater than 35Å (supplementary figure 1), which makes it difficult to infer that this mutation might have a direct action on cleavage process on S1/S2 domains. Furthermore, a recent study showed possible changes in pathogenicity derived from mutations^37^. However, the D614G variation was not observed in the sequences analyzed by this study.

#### S2 DOMAIN

Five amino acid variations (T791I, D839Y, V1176F, C1254F, and P1263L) were found in the S2 domain with frequency >1%. These variations do not happen more than one continent. Located on the HR1 region periphery (figure 3 – side in), S940F is present in Europe (0.69% of sequences) and Oceania (3.51% of sequences). T791I that have 1,09% of frequency is occurs on FP region and it is present only in Asian sequences. The V1176 variation has been found in South America with considerable frequency (6.98% of sequences). Furthermore, C1254F and P1263L variations in the IC region at the cytoplasmic end structure (figure 3 – side in) appear in Oceania (3.51% of sequences) and Europe (1.97% of sequences), respectively. These latter verified variations in IC region are residues with non-homologous physical-chemical characteristics and they might carry changes on intra-cellular portion interaction with the cellular components as well as lipid bilayer. Thus, further studies must be performed to verify the influence of these changes on SARS-CoV-2 replication.

### Epitope Variations Mapping

Our antigenic predictions show that the ELIPRO server identify the entire NTD domain as an antigen (table 2 and figure 4). In this domain, seven variations of amino acid residues were identified with non-homologous physical-chemical characteristics. The remaining 19 epitopes in the NTD had one or two variations (table 2). In contrast, the RBD domain showed only five epitopes (table 2). Recently the cryo-electron microscopy structure of the SARS-CoV-2 protein Spike trimer was determined^8,16^, showing that RBD can undergo movements between its “up” or “down” conformations. This suggest that the target epitope of the neutralizing antibody (CR3022) is accessible when the RBD is in the “up” conformation. Additionally, the ACE2 host can interact with the RBD when protein Spike is in the “up” conformation^45^. Consequently, the RBD domain has been identified as the most promising target for vaccine prototypes development^14^. However, up to now, little information about the natural variability of SARS-CoV-2 residues involved in ACE2 binding has been elucidated in the literature. Finally, the S2 domain presented 14 epitopes (three are shared with the S1 domain), two of these S2’s epitopes where found on the fusion peptide (FP) region, and another two where located on the transmembrane (TM) region.

#### T cell epitopes

Among all epitopes identified, only S^610-620^ and S^612-620^ epitopes have amino acid variation at high frequencies and worldwide prevalence (residue 614 in table1). These two epitopes have been identified by two prediction methods used in this study for T-cytotoxic cells. However, future studies are needed to find out if there is the importance of this variation in the response to the SARS-CoV-2 infection. Three more predicted epitopes for T-cytotoxic cells with variations (>1% frequency) were identified in our analyzes, two at NTD (S^50-58^ in Asia and S^240-249^ in Oceania) and a third epitope S^786-794^ in S2 domain on Asia (Highlighted in Tab 2). The prediction for T-Helper cells epitopes identified seven epitopes with variations that haven >1%frequency (Highlighted in Tab 2). The first T-Helper epitope S^50-64^ is found in Asia. The last six epitopes are co-located among S^233-253^ region in Oceania sequences (figure 4). This analysis identified the overlapping of antigenic regions for T cells in the NTD domain, but not in the RBD domain (figure 4).

Conversely, most of epitope predictions with variations >1% frequency was observed on a single continent. This suggests the existence of a random Spike protein diversification on these regions. However, the limited available literature of serological responses diversity to SARS-CoV-2 does not allow to better clarify this context at this time.

#### B cell epitopes

The total of six epitopes predicted for B cells was identified with variations at frequencies> 1%. Three linear and three conformational (highlighted in table 2). Of these, four epitopes are present in the RBD domain (only in North America), one linear (S^455-478^), and three conformational (S^434-511^, S^434-580^, and S^434-584^). The RBD polypeptide chain is classically recognized for its potential to generate neutralizing antibodies to SARS ^39,46^and recently observed also being the target of a neutralizing response for SARS-CoV-2 ^38,47^. A recent in silico study highlighted the region 524-598, which is partially present in the RBD, as one of the dominant epitopes for SARS and SARS-CoV-2 sharing an 80% identity^15^. Finally, the last two linear epitopes identified by our analysis are in domain S2 (S^786-800^, S^828-843^) with variations in Asia and Europe, respectively. Furthermore, the fusion peptide structure has been shown as a target for neutralizing immune response to SARS and SARS-CoV-2 ^16^. Variations in the S^786-800^epitope of FP region described in this study may also cause changes in its antigenicity, but, as already mentioned, new studies should be done to verify this hypothesis.

The potential variations in physicochemical characteristics brought by a amino acid residue exchange can generate an eventual condition for the viral particle escaping from the immune system via a non-neutralizing cross-reactive response from previous infections. As example, previous mutational assays with the polypeptide chain of the SARS-Cov Spike protein have shown that changes in the physical-chemical characteristics of various residues have resulted in impaired functions^5,7,48^.

Thus, there is a clear need to monitor the antigenic variation by the newly emerged coronavirus, as variations in the RBD domain may change over time, due to selective pressure on the virus through future therapies given to host populations. Although we did not identify variations in potential epitopes capable of inducing humoral response against SARS-CoV-2 with high frequencies and/or wide distribution on the continents, these slight variations covered by four months of SARS-CoV-2 spread just strengths the necessity to understand the potential of antigenic variation that the Spike protein might present. However, given the small variation in the epitopes pointed out in this work, we can suggest that vaccine approaches using the spike protein structure are unlikely to be impaired by the variability presented by theSARS-CoV-2. Still, it would be interesting to formulate therapies that focus on the S2 domain since it presented the lower number of variants. Moreover, among the S1 domain, RBD is the less affected by the variants, and therefore, more promising than NTD.

The great global engagement in tackling this pandemic has generated hundreds of vaccine prototypes and potential drugs in the short/medium term. However, prototype generation needs to rely on and be developed based on antigenic and structural variability studies, especially protein S, which is essential for virus/host interactions..

Ideally, drug and vaccine developments should take into account the virus entire diversity of its population.. Thus, efforts should focus on a continuous study of the genomic variations and their implications for the change in antigens, to guide the production of next-generation vaccines and drugs effective against all strains of SARS-CoV-2.

## Acknowledgements

This work was supported by the Conselho Nacional de Desenvolvimento Científico e Tecnológico under Grant 432611/2016-9. R.L.G., T.C.R.L., B.P.D., C.C.S.C. and C.C.B. received fellowships from Coordenação de Aperfeiçoamento de Pessoal de Nível Superior, Brasil (CAPES). A.C.G.S. received fellowships from Conselho Nacional de Desenvolvimento Científico e Tecnológico (CNPq – PIBIC).

## Declaration of interest statement

The authors declare that there is no conflict of interest.

## Author contributions

Ricardo Lemes Gonçalves: Conception of the methodology, data collection, processing, analysis and writing. Túlio César Rodrigues Leite: Data collection, processing and writing. Camila Carla da Silva Caetano: Data analysis and writing. Ana Clara Gomes de Souza, Bruna de Paula Dias, Camila Cavadas Barbosa and Ubiratan da Silva Batista: Data collection and processing. Arturo Reyes-Sandoval: Writing. Luiz Felipe Leomil Coelho: Data analysis and writing. Breno de Mello Silva: Conception of the methodology, data analysis and writing.

## Notes

### Competing Interest Statement

The authors have declared no competing interest.

