## Supplemental material 1 for "SARS-CoV-2 mutations and where to find them: An *in silico* perspective of structural changes and antigenicity of the Spike protein": supplemental material.docx

*Video 1. Structure of protein Spike of SARS-CoV2.* <https://drive.google.com/open?id=1SD1weh6YF4jSaBD4CyENfYLWNx8Td9Rg>

*Figure 1. Ramachandran plot of the three-dimensional homo-trimer model of the SARS-COV-2 spike protein.* Distribution of residues from their angles Ψ (psi) versus φ (phi). Favorable regions delimited by light blue lines and dark blue lines delimited acceptable regions. Residue outliers represented by pink circles.

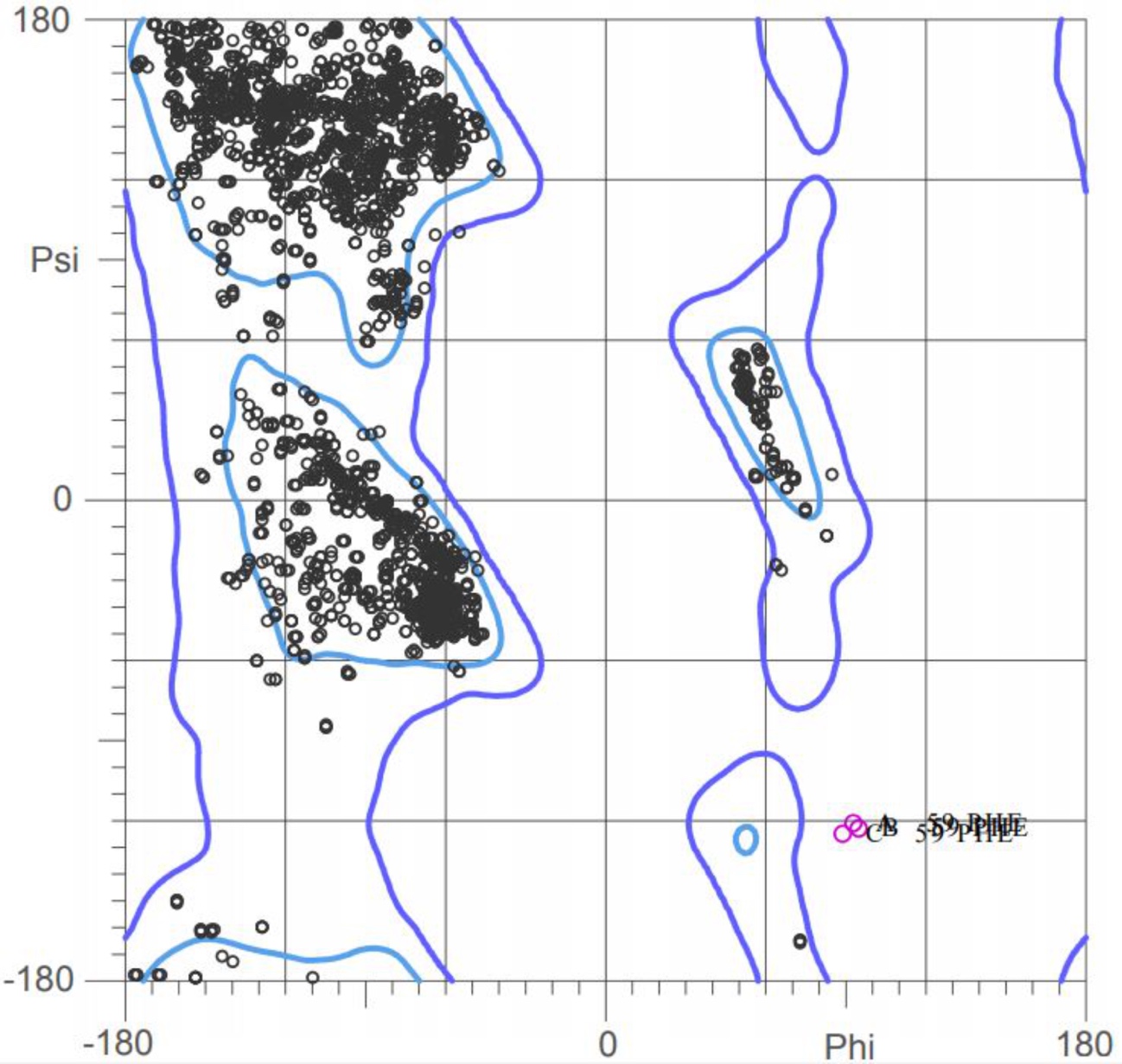

*Figure 2. The distance of alpha carbons between residues 614 and 685.* Structure of the chain A of the SARS-CoV-2 protein Spike represented in the cartoon. The distance value corresponds to angstroms.

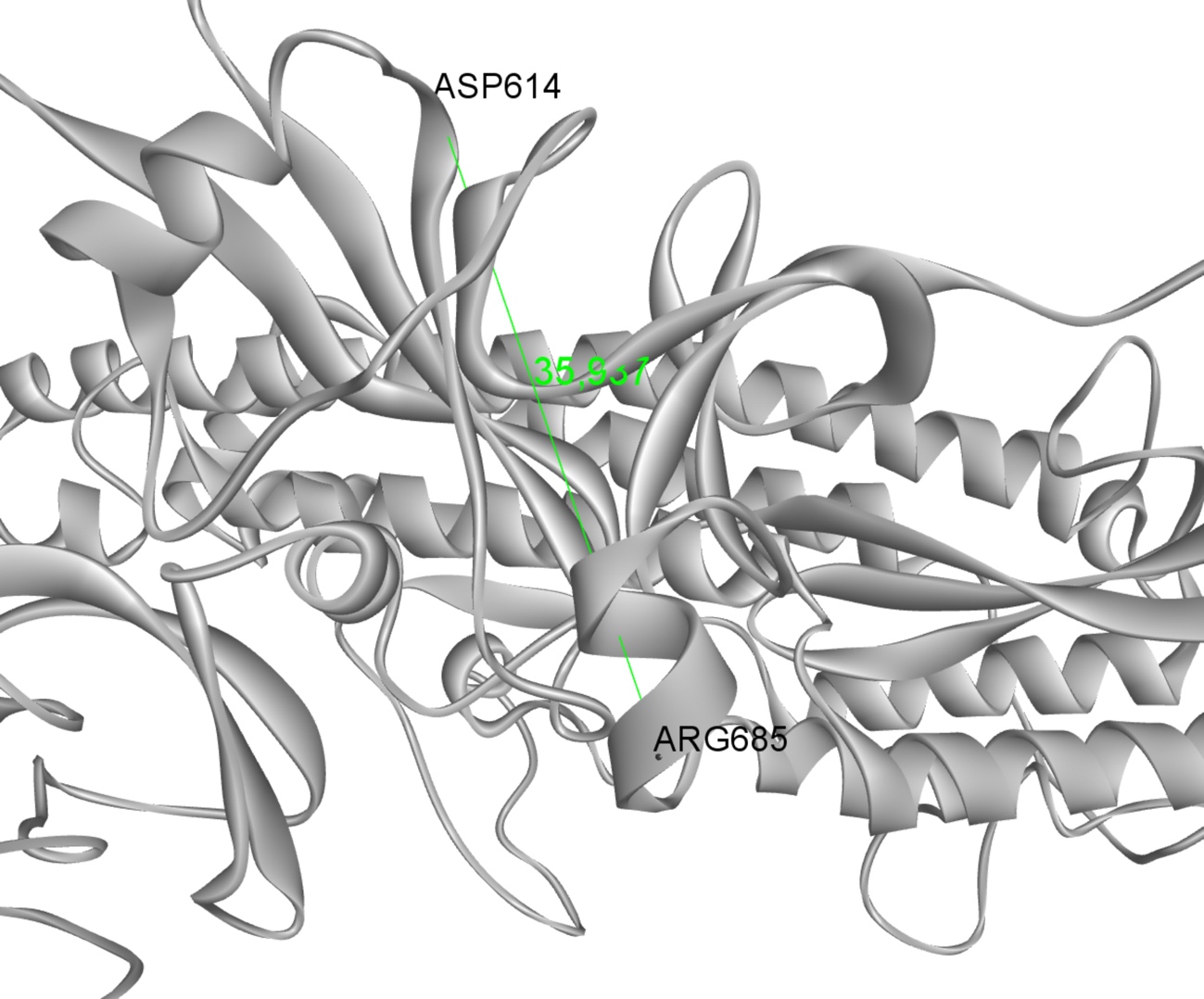

*Table 1. Overall SARS-CoV2 protein Spike epitopes.* The columns correspond, from left to right: to the forecasting methods, to the positions in the polypeptide chain in which the epitopes are located, to the peptide residues.

| **Method** | **Position** | **Peptide** |
| --- | --- | --- |
| **NetMHCpan 4.0** | 2-10 | FVFLVLLPL |
|  | 2-11 | FVFLVLLPLV |
|  | 4-11 | FLVLLPLV |
|  | 6-16 | VLLPLVSSQCV |
|  | 55-62 | FLPFFSNV |
|  | 62-70 | VTWFHAIHV |
|  | 83-93 | VLPFNDGVYFA |
|  | 109-117 | TLDSKTQSL |
|  | 117-126 | LLIVNNATNV |
|  | 133-141 | FQFCNDPFL |
|  | 133-143 | FQFCNDPFLGV |
|  | 223-233 | LEPLVDLPIGI |
|  | 240-249 | TLLALHRSYL |
|  | 268-277 | GYLQPRTFLL |
|  | 269-277 | YLQPRTFLL |
|  | 269-278 | YLQPRTFLLK |
|  | 275-285 | FLLKYNENGTI |
|  | 334-344 | NLCPFGEVFNA |
|  | 386-395 | KLNDLCFTNV |
|  | 417-425 | KIADYNYKL |
|  | 424-433 | KLPDDFTGCV |
|  | 512-520 | VLSFELLHA |
|  | 514-524 | SFELLHAPATV |
|  | 515-524 | FELLHAPATV |
|  | 610-620 | VLYQDVNCTEV |
|  | 612-620 | YQDVNCTEV |
|  | 691-699 | SIIAYTMSL |
|  | 713-722 | AIPTNFTISV |
|  | 718-726 | FTISVTTEI |
|  | 721-729 | SVTTEILPV |
|  | 786-794 | KQIYKTPPI |
|  | 817-826 | FIEDLLFNKV |
|  | 821-829 | LLFNKVTLA |
|  | 855-865 | FNGLTVLPPLL |
|  | 857-865 | GLTVLPPLL |
|  | 869-877 | MIAQYTSAL |
|  | 900-909 | MQMAYRFNGI |
|  | 921-931 | KLIANQFNSAI |
|  | 975-984 | SVLNDILSRL |
|  | 976-984 | VLNDILSRL |
|  | 983-991 | RLDKVEAEV |
|  | 999-1008 | GRLQSLQTYV |
|  | 1000-1008 | RLQSLQTYV |
|  | 1047-1056 | YHLMSFPQSA |
|  | 1048-1056 | HLMSFPQSA |
|  | 1059-1068 | GVVFLHVTYV |
|  | 1060-1068 | VVFLHVTYV |
|  | 1062-1070 | FLHVTYVPA |
|  | 1095-1104 | FVSNGTHWFV |
|  | 1192-1200 | NLNESLIDL |
|  | 1209-1218 | YIKWPWYIWL |
|  | 1215-1224 | YIWLGFIAGL |
|  | 1217-1227 | WLGFIAGLIAI |
|  | 1219-1228 | GFIAGLIAIV |
|  | 1220-1230 | FIAGLIAIVMV |
|  | 1220-1228 | FIAGLIAIV |
|  | 1220-1229 | FIAGLIAIVM |
|  | 1223-1232 | GLIAIVMVTI |
| **NetCTL 1.2** | 30-38 | NSFTRGVYY |
|  | 50-58 | STQDLFLPF |
|  | 83-91 | VLPFNDGVY |
|  | 136-144 | CNDPFLGVY |
|  | 152-160 | WMESEFRVY |
|  | 160-168 | YSSANNCTF |
|  | 162-170 | SANNCTFEY |
|  | 192-200 | FVFKNIDGY |
|  | 196-204 | NIDGYFKIY |
|  | 258-266 | WTAGAAAYY |
|  | 261-269 | GAAAYYVGY |
|  | 285-293 | ITDAVDCAL |
|  | 296-304 | LSETKCTLK |
|  | 343-351 | NATRFASVY |
|  | 357-365 | RISNCVADY |
|  | 361-369 | CVADYSVLY |
|  | 370-378 | NSASFSTFK |
|  | 372-380 | ASFSTFKCY |
|  | 392-400 | FTNVYADSF |
|  | 445-453 | VGGNYNYLY |
|  | 465-473 | ERDISTEIY |
|  | 604-612 | TSNQVAVLY |
|  | 612-620 | YQDVNCTEV |
|  | 628-636 | QLTPTWRVY |
|  | 652-660 | GAEHVNNSY |
|  | 687-695 | VASQSIIAY |
|  | 733-741 | KTSVDCTMY |
|  | 746-754 | STECSNLLL |
|  | 748-756 | ECSNLLLQY |
|  | 815-823 | RSFIEDLLF |
|  | 865-873 | LTDEMIAQY |
|  | 880-888 | GTITSGWTF |
|  | 1039-1047 | RVDFCGKGY |
|  | 1095-1103 | FVSNGTHWF |
|  | 1096-1104 | VSNGTHWFV |
|  | 1237-1245 | MTSCCSCLK |
|  | 1264-1272 | VLKGVKLHY |
| **NetMHCII 2.3** | 50-64 | STQDLFLPFFSNVTW |
|  | 51-65 | TQDLFLPFFSNVTWF |
|  | 52-66 | QDLFLPFFSNVTWFH |
|  | 53-67 | DLFLPFFSNVTWFHA |
|  | 54-68 | LFLPFFSNVTWFHAI |
|  | 165-179 | NCTFEYVSQPFLMDL |
|  | 166-180 | CTFEYVSQPFLMDLE |
|  | 167-181 | TFEYVSQPFLMDLEG |
|  | 199-213 | GYFKIYSKHTPINLV |
|  | 200-214 | YFKIYSKHTPINLVR |
|  | 201-215 | FKIYSKHTPINLVRD |
|  | 202-216 | KIYSKHTPINLVRDL |
|  | 232-246 | GINITRFQTLLALHR |
|  | 233-247 | INITRFQTLLALHRS |
|  | 234-248 | NITRFQTLLALHRSY |
|  | 235-249 | ITRFQTLLALHRSYL |
|  | 236-250 | TRFQTLLALHRSYLT |
|  | 237-251 | RFQTLLALHRSYLTP |
|  | 238-252 | FQTLLALHRSYLTPG |
|  | 239-253 | QTLLALHRSYLTPGD |
|  | 262-276 | AAAYYVGYLQPRTFL |
|  | 263-277 | AAYYVGYLQPRTFLL |
|  | 264-278 | AYYVGYLQPRTFLLK |
|  | 265-279 | YYVGYLQPRTFLLKY |
|  | 266-280 | YVGYLQPRTFLLKYN |
|  | 267-281 | VGYLQPRTFLLKYNE |
|  | 268-282 | GYLQPRTFLLKYNEN |
|  | 269-283 | YLQPRTFLLKYNENG |
|  | 374-388 | FSTFKCYGVSPTKLN |
|  | 375-389 | STFKCYGVSPTKLND |
|  | 376-390 | TFKCYGVSPTKLNDL |
|  | 509-523 | RVVVLSFELLHAPAT |
|  | 510-524 | VVVLSFELLHAPATV |
|  | 511-525 | VVLSFELLHAPATVC |
|  | 512-526 | VLSFELLHAPATVCG |
|  | 513-527 | LSFELLHAPATVCGP |
|  | 514-528 | SFELLHAPATVCGPK |
|  | 515-529 | FELLHAPATVCGPKK |
|  | 538-552 | CVNFNFNGLTGTGVL |
|  | 539-553 | VNFNFNGLTGTGVLT |
|  | 540-554 | NFNFNGLTGTGVLTE |
|  | 541-555 | FNFNGLTGTGVLTES |
|  | 542-556 | NFNGLTGTGVLTESN |
|  | 689-703 | SQSIIAYTMSLGAEN |
|  | 690-704 | QSIIAYTMSLGAENS |
|  | 691-705 | SIIAYTMSLGAENSV |
|  | 692-706 | IIAYTMSLGAENSVA |
|  | 693-707 | IAYTMSLGAENSVAY |
|  | 694-708 | AYTMSLGAENSVAYS |
|  | 695-709 | YTMSLGAENSVAYSN |
|  | 758-772 | SFCTQLNRALTGIAV |
|  | 759-773 | FCTQLNRALTGIAVE |
|  | 760-774 | CTQLNRALTGIAVEQ |
|  | 761-775 | TQLNRALTGIAVEQD |
|  | 762-776 | QLNRALTGIAVEQDK |
|  | 763-777 | LNRALTGIAVEQDKN |
|  | 850-864 | ICAQKFNGLTVLPPL |
|  | 851-865 | CAQKFNGLTVLPPLL |
|  | 852-866 | AQKFNGLTVLPPLLT |
|  | 853-867 | QKFNGLTVLPPLLTD |
|  | 854-868 | KFNGLTVLPPLLTDE |
|  | 855-869 | FNGLTVLPPLLTDEM |
|  | 865-879 | LTDEMIAQYTSALLA |
|  | 866-880 | TDEMIAQYTSALLAG |
|  | 867-881 | DEMIAQYTSALLAGT |
|  | 868-882 | EMIAQYTSALLAGTI |
|  | 869-883 | MIAQYTSALLAGTIT |
|  | 870-884 | IAQYTSALLAGTITS |
|  | 882-896 | ITSGWTFGAGAALQI |
|  | 883-897 | TSGWTFGAGAALQIP |
|  | 884-898 | SGWTFGAGAALQIPF |
|  | 885-899 | GWTFGAGAALQIPFA |
|  | 886-900 | WTFGAGAALQIPFAM |
|  | 887-901 | TFGAGAALQIPFAMQ |
|  | 888-902 | FGAGAALQIPFAMQM |
|  | 889-903 | GAGAALQIPFAMQMA |
|  | 890-904 | AGAALQIPFAMQMAY |
|  | 892-906 | AALQIPFAMQMAYRF |
|  | 893-907 | ALQIPFAMQMAYRFN |
|  | 894-908 | LQIPFAMQMAYRFNG |
|  | 895-909 | QIPFAMQMAYRFNGI |
|  | 896-910 | IPFAMQMAYRFNGIG |
|  | 897-911 | PFAMQMAYRFNGIGV |
|  | 957-971 | QALNTLVKQLSSNFG |
|  | 958-972 | ALNTLVKQLSSNFGA |
|  | 959-973 | LNTLVKQLSSNFGAI |
|  | 960-974 | NTLVKQLSSNFGAIS |
|  | 961-975 | TLVKQLSSNFGAISS |
|  | 962-976 | LVKQLSSNFGAISSV |
|  | 964-978 | KQLSSNFGAISSVLN |
|  | 965-979 | QLSSNFGAISSVLND |
|  | 966-980 | LSSNFGAISSVLNDI |
|  | 967-981 | SSNFGAISSVLNDIL |
|  | 968-982 | SNFGAISSVLNDILS |
|  | 969-983 | NFGAISSVLNDILSR |
|  | 989-1003 | AEVQIDRLITGRLQS |
|  | 990-1004 | EVQIDRLITGRLQSL |
|  | 991-1005 | VQIDRLITGRLQSLQ |
|  | 992-1006 | QIDRLITGRLQSLQT |
|  | 993-1007 | IDRLITGRLQSLQTY |
|  | 994-1008 | DRLITGRLQSLQTYV |
|  | 995-1009 | RLITGRLQSLQTYVT |
|  | 996-1010 | LITGRLQSLQTYVTQ |
|  | 997-1011 | ITGRLQSLQTYVTQQ |
|  | 998-1012 | TGRLQSLQTYVTQQL |
|  | 999-1013 | GRLQSLQTYVTQQLI |
|  | 1000-1014 | RLQSLQTYVTQQLIR |
|  | 1001-1015 | LQSLQTYVTQQLIRA |
|  | 1002-1016 | QSLQTYVTQQLIRAA |
|  | 1003-1017 | SLQTYVTQQLIRAAE |
|  | 1004-1018 | LQTYVTQQLIRAAEI |
|  | 1005-1019 | QTYVTQQLIRAAEIR |
|  | 1006-1020 | TYVTQQLIRAAEIRA |
|  | 1007-1021 | YVTQQLIRAAEIRAS |
|  | 1008-1022 | VTQQLIRAAEIRASA |
|  | 1009-1023 | TQQLIRAAEIRASAN |
|  | 1010-1024 | QQLIRAAEIRASANL |
|  | 1014-1028 | RAAEIRASANLAATK |
|  | 1015-1029 | AAEIRASANLAATKM |
|  | 1041-1055 | DFCGKGYHLMSFPQS |
|  | 1042-1056 | FCGKGYHLMSFPQSA |
|  | 1043-1057 | CGKGYHLMSFPQSAP |
|  | 1044-1058 | GKGYHLMSFPQSAPH |
|  | 1045-1059 | KGYHLMSFPQSAPHG |
|  | 1046-1060 | GYHLMSFPQSAPHGV |
|  | 1047-1061 | YHLMSFPQSAPHGVV |
|  | 1057-1071 | PHGVVFLHVTYVPAQ |
|  | 1058-1072 | HGVVFLHVTYVPAQE |
|  | 1059-1073 | GVVFLHVTYVPAQEK |
|  | 1070-1084 | AQEKNFTTAPAICHD |
|  | 1071-1085 | QEKNFTTAPAICHDG |
|  | 1072-1086 | EKNFTTAPAICHDGK |
|  | 1073-1087 | KNFTTAPAICHDGKA |
|  | 1099-1113 | GTHWFVTQRNFYEPQ |
|  | 1100-1114 | THWFVTQRNFYEPQI |
| **BepiPred-2.0** | 14-33 | QCVNLTTRTQLPPAYTNSFT |
|  | 59-81 | FSNVTWFHAIHVSGTNGTKRFDN |
|  | 141-163 | LGVYYHKNNKSWMESEFRVYSSA |
|  | 178-191 | DLEGKQGNFKNLRE |
|  | 207-222 | HTPINLVRDLPQGFSA |
|  | 248-260 | YLTPGDSSSGWTA |
|  | 313-322 | YQTSNFRVQP |
|  | 331-336 | NITNLC |
|  | 338-356 | FGEVFNATRFASVYAWNRK |
|  | 370-395 | NSASFSTFKCYGVSPTKLNDLCFTNV |
|  | 404-424 | GDEVRQIAPGQTGKIADYNYK |
|  | 439-451 | NNLDSKVGGNYNYL |
|  | 455-478 | LFRKSNLKPFERDISTEIYQAGST |
|  | 483-493 | VEGFNCYFPLQ |
|  | 497-502 | FQPTNG |
|  | 516-535 | ELLHAPATVGCPKKSTNLVK |
|  | 615-621 | VNCTEVP |
|  | 626-645 | ADQLTPTWRVYSTGSNVFQT |
|  | 656-665 | VNNSYECDIP |
|  | 673-691 | SYQTQTNSPRRARSVASQS |
|  | 694-709 | AYTMSLGAENSVAYSN |
|  | 786-800 | KQIYKTPPIKDFGGF |
|  | 806-815 | LPDPSKPSKR |
|  | 828-843 | LADAGFIKQYGDCLGD |
|  | 987-993 | VEAEVQI |
|  | 1035-1043 | GQSKRVDFC |
|  | 1109-1118 | FYEPQIITTD |
|  | 1133-1172 | VNNTVYDPLQPELDSFKEELDKYFKNHTSP |
|  |  | DVDLGDISGI |
|  | 1202-1207 | ELGKYE |
|  | 1252-1270 | SCCKFDEDDSEPVLKGVKL |
| **ElliPro** | 14-38 | QCVNLTTRTQLPPAYTNSFTRGVYY |
|  | 56-270 | QCVNLTTRTQLPPAYTNSFTRGVYYPDKVFR |
|  |  | SSVLHSTQDLFLPFFSNVTWFHAIHVSGTNG |
|  |  | TKRFDNPVLPFNDGVYFASTEKSNIIRGWIFG |
|  |  | TTLDSKTQSLLIVNNATNVVIKVCEFQFCNDP |
|  |  | FLGVYYHKNNKSWMESEFRVYSSANNCTFE |
|  |  | YVSQPFLMDLEGKQGNFKNLREFVFKNIDG |
|  |  | YFKIYSKHTPINLVRDLPQGFSALEPLVDLPIG |
|  |  | INITRFQTLLALHRSYLTPGDSSSGWTAGAAA |
|  |  | YYVGYL |
|  | 323-375 | TESIVRFPNITNLCPFGEVFNATRFASVYAWN |
|  |  | RKRISNCVADYSVLYNSASFS |
|  | 391-404 | CFTNVYADSFVIRG |
|  | 391-424 | CFTNVYADSFVIRGGKYNYK |
|  | 434-511 | IAWNSNNLDSKVGGNYNYLYRLFRKSNLKPFE |
|  |  | RDISTEIYQAGSTPCNGVEGFNCYFPLQSYGFQ |
|  |  | PTNGVGYQPYRVV |
|  | 434-580 | IAWNSNNLDSKVGGNYNYLYRLFRKSNLKPFE |
|  |  | RDISTEIYQAGSTPCNGVEGFNCYFPLQSYGFQ |
|  |  | PTNGVGYQPYRVVVVLSFELLHAPATVCGPKKS |
|  |  | TNLVKNKCVNFNFNGLTGTGVLTESNKKFLPFQ |
|  |  | QFGRDIADTTDAVRDPQ |
|  | 434-584 | IAW...TLEI |
|  | 544-564 | LHAPATVCGPKKSTNLVKNK |
|  | 577-584 | ESNKFLPFQQRDPQTLEI |
|  | 636-641 | YSTGSN |
|  | 675-690 | QTQTNSPRRARSVASQ |
|  | 675-691 | QTQTNSPRRARSVASQS |
|  | 1078-1088 | APAICHDGKAH |
|  | 1096-1103 | VSNGTHWF |
|  | 1110-1116 | YEPQIIT |
| **Discotope-2.0** | 443-449 | SKVGGNY |
|  | 496-505 | GFQPTNGVGY |
|  | 681-688 | PRRARSVA |
|  | 1138-1160 | YDPLQPELDSFKEELDKYFKNHT |
